# Co-adaption of tRNA Gene Copy Number and Amino Acid Usage Influences Translation Rates in Three Life Domains

**DOI:** 10.1101/104570

**Authors:** Meng-Ze Du, Wen Wei, Lei Qin, Shuo Liu, An-Ying Zhang, Yong Zhang, Hong Zhou, Feng-Biao Guo

## Abstract

The cellular translation process should obey the principle of maximizing efficiency and minimizing resource and energy costs. Here, we validated this principle by focusing on the basic translation components of tRNAs and amino acids. To most efficiently utilize these components, we reasoned that the quantities of the 20 tRNAs and their corresponding amino acids would be consistent in an organism. The two values should match at both the organismal and protein scales. For the former, they co-vary to meet the need to translate more proteins in fast-growing or larger cells. For the latter, they are consistent to different extents for various proteins in an organism to comply with different needs of translation speed. In this work, 310 out of 410 genomes in three domains had significant co-adaptions between the tRNA gene copy number and amino acid composition, and thus validating the principle at the organism scale. Furthermore, fast-growing bacteria co-adapt better than slow-growing ones. Highly expressed proteins and those connected to acute responses have better co-adaption, illustrating the principle at the individual protein scale. Experimentally, manipulating the tRNA gene copy number to optimize co-adaption between enhanced green fluorescent protein (EGFP) and tRNA gene set of *Escherichia coli* indeed lifted the translation rate (speed). Our results also contribute to revealing a translation rate-associated factor with universal and global effects. From a practical perspective, our findings suggest a strategy to increase the expression of target proteins and have implications for designing chassis cells in the field of synthetic biology field.

## Introduction

Translation initiation, elongation and termination involve many factors, that balance translation rate (speed) and accuracy (Gingold and Pilpel 2011; Yang et al. 2014). The final translation efficiency for a given protein is restricted by the cost of its production and organization (Dekel and Alon 2005). Therefore, evolving a genome-wide translation regulation regime can efficiently determine the translation rates of various genes in different conditions (Gingold and Pilpel 2011). Conventional computations of translation elongation efficiency refer to codon usage (Sharp and Li 1987) and tRNA availability (dos Reis et al. 2004). The relationship between codon usage and tRNA abundance predicts translation efficiency with reasonable accuracy (Gingold and Pilpel 2011).

Additional theories have been proposed with constantly emerging experimental technologies (Ingolia 2016; Yan et al. 2016) to cope with challenges to the simplified assumptions about translation described above. Thus, the effects of codon order (Fredrick and Ibba 2010; Tuller et al. 2010a; Gamble et al. 2016), local tRNA availability (Elf et al. 2003; Chan and Lowe 2009; Nedialkova and Leidel 2015), regulation of expression of the tRNA gene (Cannarozzi et al. 2010), the diverse demands of the transcriptomes (Dittmar et al. 2006; Tuller et al. 2010a), ribosomes (Qu et al. 2011), mRNA structures (Li et al. 2012; Shah et al. 2013; Yu et al. 2015) and folding energy (Tuller et al. 2010b) were included in the translation efficiency models. Among these factors, tRNA availability repeatedly emphasized decides the supply of aminoacyl-tRNA (Vargas-Rodriguez and Musier-Forsyth 2014), which influences the translocation of ribosomes on mRNA (Subramaniam et al. 2014; Espah Borujeni and Salis 2016; Rozov et al. 2016; Wu et al. 2016). Nutriment limitations, such as amino acid shortage, also have influences on the cellular supply of aminoacyl-tRNA (Mandel and Silhavy 2005). However, how the formation of aminoacyl-tRNA influences translation efficiency is still unclear.

In the translation process, tRNAs can be thought of as tools and the amino acids as the raw materials. Each species of tRNA corresponds to a particular amino acid, and each of the former is responsible for carrying one of the latter. We hypothesize that the levels of the tRNAs and the corresponding amino acids should be well matched to synthesize proteins more efficiently. Such a consistency would maximize efficiency and minimize resource/energy costs.

Here, we try to test the concept that the process of translation is selected for maximum efficiency by examining the association between the tRNA gene copy numbers and amino acid compositions in various organisms. We sought to validate two points of reasoning: at the organismal scale, most organisms evolve co-adaption between tRNA gene copy number and amino acid composition; and at the second scale of individual proteins, the co-adaption intensity may vary among the proteins within an organism. Some proteins need to be expressed rapidly to maintain a high quantity or to satisfy the requirements of acute responses. We speculated that such proteins would have higher co-adaption to increase their translation efficiency Computational analyses were employed to elucidate co-adaption between tRNA gene copy numbers and amino acid usage for proteins/proteomes in three domains of life, indicating the effects of maximum efficiency and the minimum cost principle. Then, we correlated the co-adaption with proteins’ translation rates, which were validated by growth rates of bacteria and production rates of target proteins. The target proteins’ translation rates were observed to be lifted through changing the proportions of gene copy numbers of tRNAs, providing clues for applied biology.

## Results

### Validation of the principle at the genome scale

During translation, tRNAs transport amino acids to ribosomes; co-operation between these factors has been reported in a few organisms (Yamao et al. 1991). Previous researches demonstrated that the tRNA gene copy numbers were different among organisms/strains/species, and that the protein sequences varied greatly (Levitt 2009). To check whether the amino acid usage of proteins generally co-adapts with the corresponding organism’s tRNA gene copy number, we calculated and compared independent frequencies of the two in 410 genomes from three domains of life (17 archaea; 359 bacteria; 34 eukaryotes), using more accurate tRNA gene annotations in GenBank (Benson et al. 2013), the Genomic tRNA database (Chan and Lowe 2009) and tRNAdb (Jühling et al. 2009). First, the frequencies of 20 standard tRNA genes (Table S1) were computed by counting cognate tRNA gene copies and being divided by the organism’s total tRNA gene counts. Second, frequencies of the 20 amino acids in the proteome (Table S1) were computed by dividing the count of each amino acid by the sum of the twenty amino acid counts. After obtaining these two types of data for each organism, we performed linear fit and correlation analyses.

The linear fit results showed variable co-adaption relationships (Fig. 1A and Table S1), illustrating that the two factors (tRNA gene copy number and the amino acid frequency) are not independent from each other. Indeed, although the slopes of the fitted lines differed, in all cases, the tRNA gene copy numbers showed positive correlations with corresponding amino acid usages. Spearman rank correlation coefficients (r) were calculated after least square fitting (Table S1). Specifically, 99.27% had correlation coefficients greater than 0.1 (Fig. 1B), and 75.61% showed significant correlations (p < 0.05). Finally, a general linear relationship exists between tRNA gene copy and amino acid usage. For four representative organisms, the archaebacterium *Methanosphaera stadtmanae* (r = 0.17, p = 0.46), the bacterium *Escherichia coli* (r = 0.54, p = 0.01), and the eukaryotes *Saccharomyces cerevisiae* (r = 0.74, p = 1.67E-04) and *Homo sapiens* (r = 0.56, p = 0.01), the linear models presented different co-adaption intensities (Fig. 1C, Table S1). Compared with the other three organisms, yeast had the best linear fit. However, for *M. stadtmanae*, having the most unbalanced constitution of tRNA genes, the observed points were not well fitted. In general, most organisms’ tRNA gene copy numbers and amino acid usages showed a positive linear relationship.

**Fig. 1.**
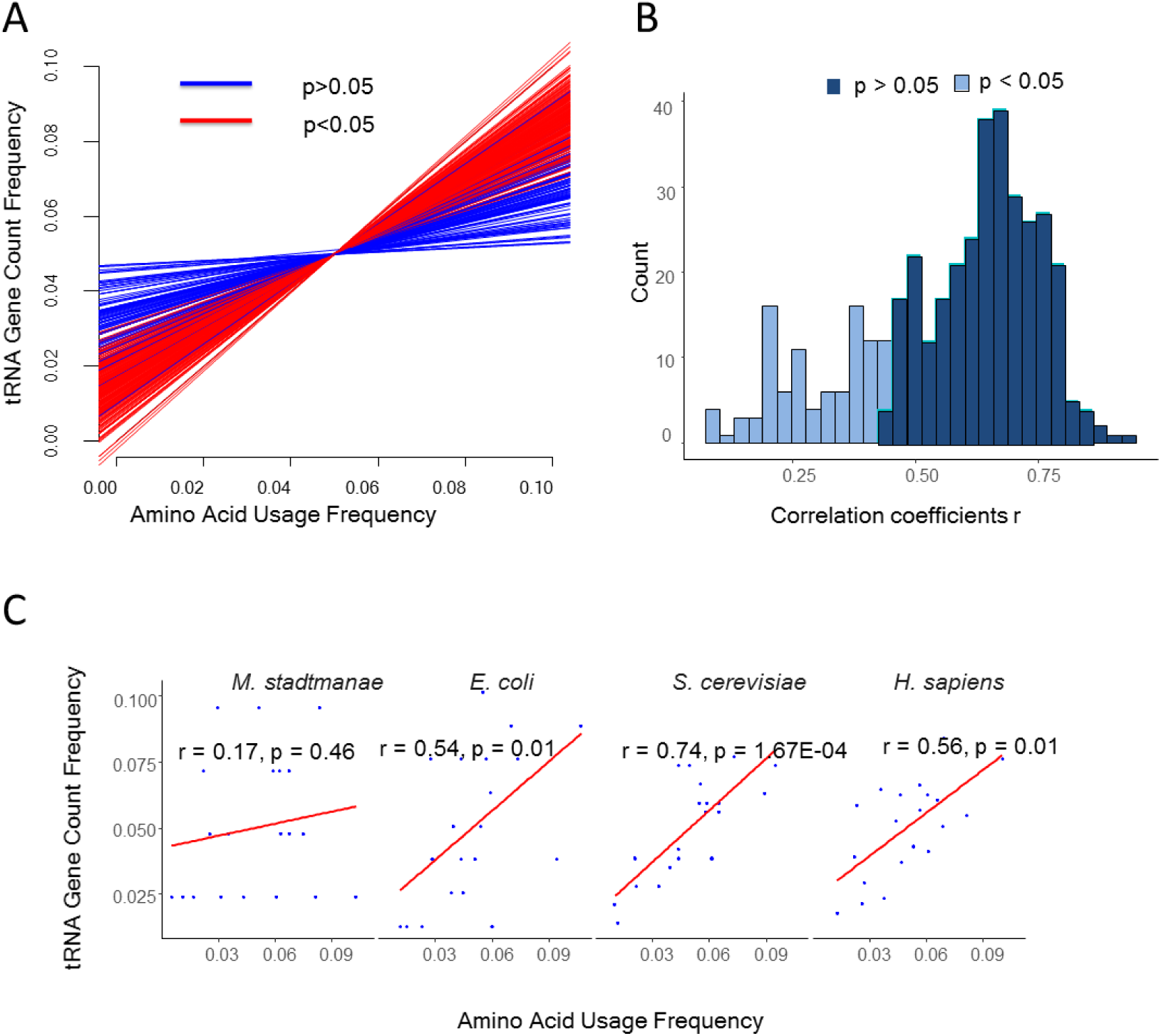
Co-adaption between frequencies of tRNA gene copy numbers and amino acid usage. A) Linear fitting results for 410 organisms. When the p value of linear fitting is greater than 0.05, the lines are blue; red indicates a p value less than 0.05. B) Corresponding Spearman rank correlation coefficients r for the linear fit of 410 genomes. C) Linear fit of four model organisms.

**Table 1.** Analaysis Of The Potentially Rapidly Expressed Protiens According To Their Functions In *E.Coil* and *S.Cerevisiae*.

|  | *E. coli* |  |  |  | *S. cerevisiae* |  |  |  |
| --- | --- | --- | --- | --- | --- | --- | --- | --- |
|  | Number | Abundance | TAAI | CAI | Number | Abundance | TAAI | CAI |
| Whole genome | 3133 | 319.2 | 0.51 | 0.35 | 6087 | 163.8 | 0.62 | 0.18 |
| Ribosome subunits | 57 | 4724.28 | 0.59 | 0.63 | 178 | 1231.65 | 0.74 | 0.48 |
| Cell division | 19 | **131.84** | 0.57 | 0.35 | 20 | **15.09** | 0.67 | **0.16** |
| Two-component system | 58 | **80.7** | 0.57 | **0.29** | 216 | 203.86 | 0.65 | 0.19 |
| Mismatch repair | 20 | **57.21** | 0.53 | **0.34** | 19 | **32.51** | 0.72 | **0.16** |
| Sugar metabolism | 47 | 325.52 | 0.52 | 0.41 | 28 | 563.53 | 0.69 | 0.26 |

Proteins in bold face had lower corresponding values than genome's average values.

We measured the co-adaption intensity using a ***t***RNA gene copy and ***a***mino ***a***cid usage accordance index (TAAI), which is close to the correlation coefficient of linear fitting and equal to the r value of the Spearman rank correlation. During protein production, tRNA genes will be transcribed to tRNAs, and then loaded with amino acids for protein translation. Resource allocation would be the most efficient if the supply of tRNAs just meets the required amount of amino acids. Based on the results, our species/strain/organism scale reasoning was confirmed. In other words, most genomes had significant co-adaption between the tRNA gene copy number and the frequencies of amino acid usage and hence maximized their translation efficiency and minimized their energy/resource costs.

Different genomes (species/strain/organism) may have different translation selection pressure: to translate different numbers of proteins in a given time. For example, large genomes have more proteins, and fast-growing bacteria need to synthesize more proteins simultaneously. In fact, large bacterial genomes are often associated with short generation times (Rocha 2004). According to the maximum efficiency/minimum cost principle, fast-growing/large bacteria should have higher TAAIs than slow-growing and/or small bacteria. To test this possibility, we compared the TAAIs of 53 bacteria (Table S2) and grouped them by growth time (Rocha 2004). The fast, had growth times below the mean of the 53 ones; the slow, had growth times greater than the average. The two groups had similar variances of TAAI values, while the slow group had significantly lower TAAIs than the fast group (Fig. 2A). Thus, co-adaption showed an effect on growth rate. This result is consistent with the idea that population growth rate is a fundamental ecological and evolutionary characteristic of living organisms (Kempes et al. 2012). Similarly, larger bacteria have larger genomes and more proteins that need to be translated than bacteria with smaller genomes (Kempes et al. 2012; Kempes et al. 2016). Therefore, we also compared the TAAIs of prokaryotic organisms grouped by genome size (small, medium and large). These three groups had significantly divergent mean TAAIs of 0.37, 0.60 and 0.65 (p value of Student’s t test: < 2.2e-16; Fig. 2B). That the relatively larger genomes have higher TAAIs supports the conclusion that bacteria under higher selective pressure have higher TAAIs, thus conforming to our first hypothesis based on the principle of efficiency described above. Bacteria with smaller genomes and slow growing speeds would suffer less pressures from protein translations and hence have lower TAAIs.

**Fig. 2.**
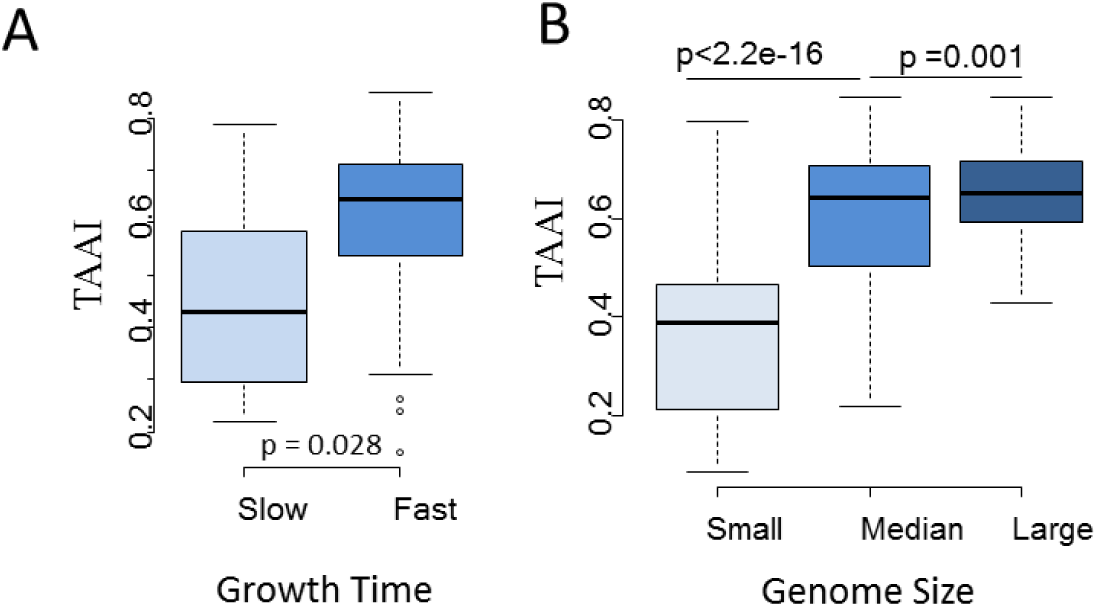
Co-adaption at the genome scale. A) TAAIs of bacteria divided into two groups based on their growth rates. The dataset includes 53 bacteria with available information on growth rates. The fast group has higher TAAIs than the slow. B) Prokaryotic organisms’ TAAIs, associated with corresponding genome sizes. The prokaryotic organisms were divided into three groups, showing significantly different TAAIs. Here, 376 prokaryotic genomes were involved in the analysis of correlation between TAAIs and genome sizes.

Some genomes have non-significant TAAIs and it would be beneficial to understand why. In prokaryotes, genome size and TAAI correlated well (r = 0.49, p < 2.2e-16) and almost all the 96 prokaryotes with bad TAAI/co-adaption do have genome sizes smaller than 2.5Mb, whereas almost all genomes with good co-adaption are larger than 2.6 Mb. The weak TAAI of smaller genomes is obviously caused by their deficiency in request of translation efficiency: less selection pressure, which could be measured based on the genome size in these organisms. However, in eukaryotes, alternative splicing of messenger RNA results in an inconsistency between the number of proteins produced and genome size. Using the quotient of the number of proteins divided by genome size should be a more reliable reflection of eukaryotes’ actual translation demand (also selection pressure). Consequently, four eukaryotes, *Bos taurus* (cow), *Felis catus* (cat), *Strongylocentrotus purpuratus* (Sea urchin) and *Plasmodium falciparum* with non-significant TAAIs indeed have smaller quotient. Therefore, it is reasonable that the lower translation demand (also selective pressure) leads some genomes to have bad TAAIs.

### Validation of the principle at the protein scale

The aforementioned results validated the principle at the genome (species/strain/organism) scale. Next, we asked whether there are co-adaption divergences within genomes and what such divergences may signify. The proteins within a genome also have different adaptions (variable TAAIs) for their different amino acid compositions (Fig. 3A). Taking *E. coli* as an example, a distinct difference was noted when comparing the co-adaption of the five proteins with the highest TAAIs and the five proteins with the lowest TAAIs (Fig. 3B). The amino acid frequencies of the top five were more consistent with the corresponding genomic tRNA gene copy frequencies. Within a given genome, this co-adaption divergence generally occurred among proteins.

**Fig. 3.**
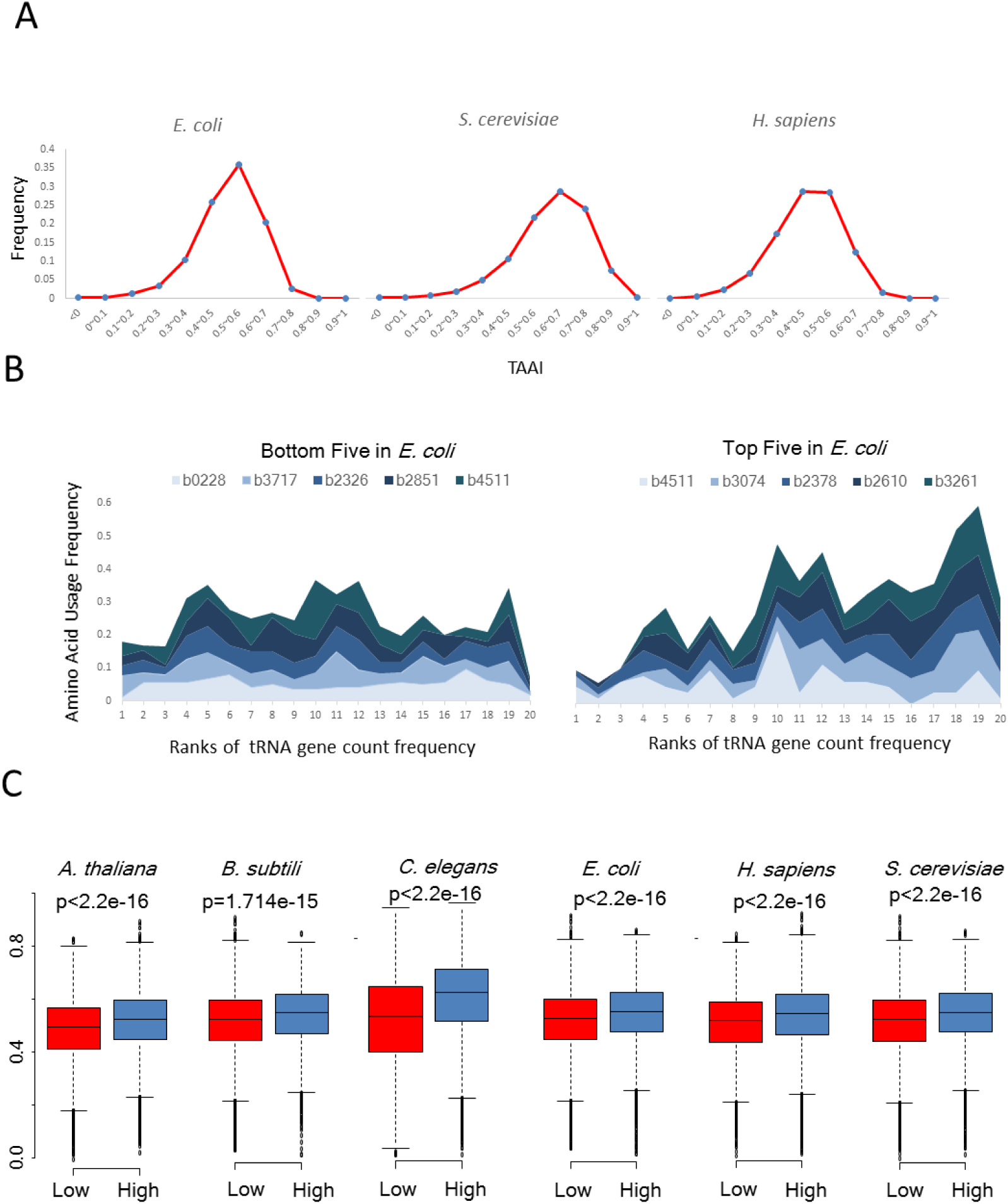
Intra-genome variation and translation factors. A) TAAI distribution in three model organisms. B) A stacked area map of the five E. coli genes with the lowest TAAI values, and a stacked area map of the five E. coli genes with the highest TAAI values. The amino acid frequencies were sorted by tRNA gene copy number. The stacked area that increases with the order of tRNA gene copy frequencies (horizontal axis) shows better co-adaption. C) Analysis o f six model organisms’ abundance shows that highly expressed proteins generally have higher TAAIs than proteins with lower expression levels. The high and low values were normalized to each genome’s median value of abundance.

Selective pressure within genomes is reflected by the direct results of translation efficiency: proteins’ abundances, even though the two values are not entirely equivalent. Co-adaption should reflect the supply of aminoacyl-tRNA, which ultimately affects the final protein synthesis. We compared six model organisms’ TAAIs and found that proteins with higher expression levels had clearly higher TAAIs than proteins with lower expression levels by Student's t test (Fig. 3C). Furthermore, when using a linear fit, the TAAIs and the protein abundances showed significantly positive correlations (p < 1e-6; Table S3). This result is consistent with the idea that tRNA level has direct effects on translation efficiency (Dana and Tuller 2014). Thus, as a reflection of the translation rate, protein abundances correlate with TAAIs to a certain extent, and their relationship seems to be a consequence of selection pressure to have a suitable translation rate.

To further explore this finding, and considering that paralogous genes in the same family have similar molecular evolutionary stresses and changes (Jost et al. 2008), we compared the TAAIs in *E. coli* and yeast according to gene function groups: ribosome subunits, cell division (Hale and de Boer 1997), two-component system (including response regulators and sensors; (Chang and Stewart 1998), mismatch repair (Kunkel and Erie 2005), and sugar metabolism (Titgemeyer and Hillen 2002). The average TAAIs and protein abundances for five groups of genes were calculated (Table 1). For the *E. coli* and yeast genomes, proteins from all groups had average TAAIs higher than the genome average (Student’s t test: p = 3.54e-12). Three of the five groups correspond to acute responses (cell division, two-component system, and mismatch repair), and the other two groups relate to fast growth (ribosome subunits and sugar metabolism). Ribosome subunits are important participants in the translation process, and there is no doubt that ribosome subunits have the highest abundance, TAAI and codon usage bias index CAI (Sharp and Li 1987). Sugar metabolism proteins, which includes proteins in “Amino sugar and nucleotide sugar metabolism”, also has higher TAAI and abundance than genome average. Although the abundances of proteins in the other three categories are lower than genome average, their TAAIs were higher. We further compared the protein abundance and TAAIs of experimentally determined upregulated yeast genes (Ingolia et al. 2009), and observed similar results (Table S4). Thus, proteins involved in acute responses under more selective pressure, generally have good co-adaption relationships.

Therefore, our second reasoning was also validated: at the scale of individual proteins, co-adaption intensity may vary among the protein collective within a genome. Some proteins need to be expressed rapidly to maintain their quantity to be connected with an acute response.

### Experimental Verification of Co-Adaption Affecting the Translation Rate

According to the principle and the above results, co-adaption was associated with the translation rate, prompting us to examine whether the proteins with high TAAIs indeed have high translation rates *in vivo*. To test this conjecture, one copy of a specific tRNA gene that might increase or decrease the TAAI was introduced to *E. coli* along with a gene encoding enhanced green fluorescent protein (EGFP) after which EGFP protein synthesis was analyzed.

Proteins with higher TAAIs might have higher translation rates, and thus higher production levels. EGFP is easy to express and detect, and constructs for tRNA overexpression have previously been designed and tested (Acosta-Rivero et al. 2002). Therefore, by combining the expression of EGFP with a tRNA gene, we can see the effect of a specific tRNA on EGFP expression. Increasing the copy number of the corresponding tRNA gene may increase or decrease the TAAI for EGFP and the whole genome (Fig. 4A). EGFP had a TAAI of 0.45 when expressed with the original frequencies of *E. coli* tRNA genes. When one gene copy encoding either 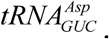, 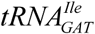 or 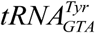 was introduced, the corresponding Δ*TAAIs* for EGFP were 0.03, 0.0008 or −0.007, respectively, and the cumulative Δ*TAAIs* for all proteins of the genome were −28.68, −43.77 or 78.68, respectively. The following sequences were constructed in plasmids: *EGFP* (control group), 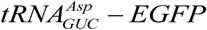, 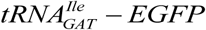, and 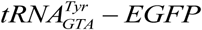 (tRNA-Tyr), which were used to transfer *E. coli* Top10 cells (Fig. 4B). In *E. coli*, there is only one type of tRNA (isoacceptor tRNA) for Asp, Tyr and Ile, which means that codon bias has no observable effect on the increase in EGFP expression. We observed fluorescence intensity with confocal microscopy and found that the EGFP yield of the experimental group expressing 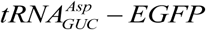 was significantly higher than the others at the same time point (Fig. 4B and Fig. S1A). Detailed fluorescence intensities of the four groups acquired with a fluorospectro photometer (Fig. 4C), showed that the EGFP production efficiency of the four groups from low to high was as follows: Control, tRNA-Ile, tRNA-Tyr and tRNA-Asp. The fluorescence intensities were consistent with western blotting results (Fig. 4D and Fig. S1B). The experimental group tRNA-Asp produced ten times more EGFP than the Control. Considering the dynamic processes involved in EGFP abundance variation, the slope of 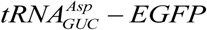 abundance was much higher than that of the control, indicating that the translation rate of the former was higher.

**Fig. 4.**
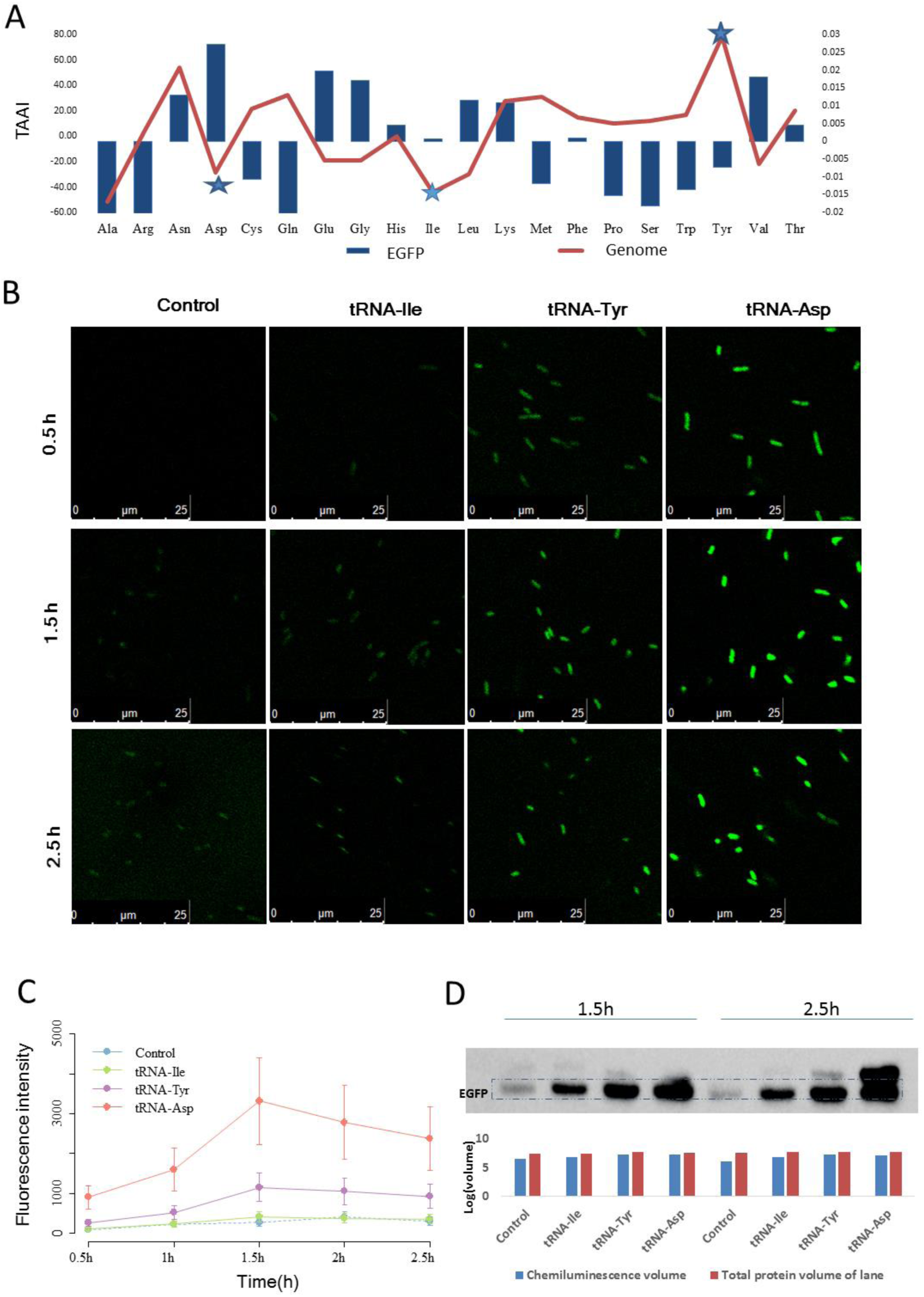
EGFP expression of with original and optimized TAAIs in *E. coli*. A) Δ*TAAI* is the upregulated TAAI value resulting from adding a copy of one of the 20 standard tRNAs. The star marks tRNA-Asp, tRNA-Ile and tRNA-Tyr. B) Confocal micrographs of the control and experimental groups present different fluorescence intensities in the EGFP channel; the corresponding merge figures of the bright-field and EGFP channel are shown in Fig. S1. C) Fluorescence intensities of four nascent sequences with EGFP at 513nm from 0 h to 2.5 h. All of these results showed that in the experimental group transformed with the Asp tRNA gene, there was an approximately ten-fold lift. D) Western blot results for the nascent sequences. The following histograms show the normalized density of the corresponding lane, and the chemiluminescence intensity of the corresponding target band using stain-free technology (Bio-Rad). The corresponding electrophoretogram, shown in Supplementary Fig. 1B, reflects the loading volume of the total proteins.

To further rule out the possibility of codon usage influence, we compared the number and coding order of codons for the three tRNAs in EGFP (Fig. S1C). The EGFP mRNA sequence has one type of Ile codon (ATC), two types of Tyr codons (TAT: 9%, TAC: 91%), and two Asp codons (GAT: 11%; GAC: 89%). All preferred codons are the corresponding codons for the *E. coli* cognate tRNAs. Therefore, there should be no significant variant effect of varying from the preferred codon. Then, we calculated the dispersion degree by analysis of variance, as the order of tRNA can influence its recycling during translation (Cannarozzi et al. 2010). We analyzed the variance of amino acid sites both locally and globally. The variance of the first eleven sites for Ile, Tyr and Asp are: 53, 61 and 46. The corresponding recycling effects should be weakened when increasing specific tRNA gene copy numbers. In fact, increasing the Ile tRNA gene copy number does not significantly increase EGFP production. Together, the results do not indicate an influence of tRNA recycling or preferred codon usage. This experiment confirmed that co-adaption has a clear effect on translation rate. Thus, optimizing co-adaption could significantly promote translation production of foreign proteins.

## Discussion

Cells are believed to evolve to maximize efficiency and minimize resource and energy cost (Maitra and Dill 2015; Grosskopf et al. 2016). We hypothesized that this principle would affect translation mechanisms, and we tested this conjecture based on the basic translation “tool” tRNAs and the “raw material” amino acids. To maximally utilize the resources, we reasoned that the quantities of the 20 tRNAs and amino acids in a species should be consistent based on this principle. For simple and convenient analysis, we used the tRNA gene copy number as the proxy for the former, and used the amino acid frequency as the latter. The genome has an average vector of amino acid frequency and each protein also has its vector form of amino acid frequency. Hence, there would be a general co-adaption value for each genome and a specific co-adaption value for each protein. Using correlation and abundance (functional group) analyses we validated our two conjectures, which are logical outcomes of the maximum efficiency and minimum cost principle. Based on the results and analyses, the genome’s TAAI could be regarded as a proxy for general translation efficiency or actual translation needs. Proteins’ TAAIs reflect the highest translation efficiency or translation need in extreme conditions.

Co-adaption is a global effect exerted on proteins and organisms. Each organism has a specific amino acid usage and a co-adapted tRNA gene copy number. This co-adaption maximizes the translation efficiency of the complete proteomes. The larger translation pressure the organism is exposed to, the higher average TAAI it has. In a genome, almost all the proteins have positive TAAI values (Fig. 3A). Hence, this co-adaption as a translation rate associated factor is applicable to all three domains of life and all proteins within an organism. In contrast to the consistency between the effects of synonymous codon usage and tRNA gene copy number on translation rates (Duret 2000; Li et al. 2012), which is a local factor acting on regions of a gene, the TAAI is a translation rate associated factors with universal and global effects.

Co-adaption arises from energy efficiency and selective pressure. Organisms evolve to maximize efficiency and minimize energy cost by adapting through genetic mechanisms (Maitra and Dill 2015; Kempes et al. 2016). Such global co-adaption might raise the translation efficiency globally, coinciding with the energy efficiency/ecological dynamics principle (Grosskopf et al. 2016). Previously, Higgs and Ran analyzed 80 bacterial genomes and found that tRNA gene copy numbers evolved in response to translational selection (Higgs and Ran 2008). It is notable that they observed consistency between synonymous codon usage and tRNA gene copy number and that the unequal usage of synonymous codons encoding the same amino acid was involved. However, our current study on co-adaption focuses on disequilibrium frequencies among the twenty standard amino acids. Here, the co-adaption reflects a balance between tRNA gene copy number and the amino acids needed by the proteome. Redundant excessive tRNA gene copies will ultimately be a waste of translation resources (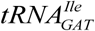 did not increase the EGFP production in *E. coli*). Selective pressure drives the co-adaption at the protein level. Experiments showed that increasing the TAAI could indeed improve the translation speed of the proteins and hence validate that the co-adaption is caused directly by translation pressure. These results indicate that translation selection causes co-variation at the scale of organisms and individual proteins.

If we expand our view to the domains of life, which have evolutionary connections (Lynch and Conery 2003; Booth et al. 2016), such selection also exists. We found that eukaryotic genomes had much better adaption values than the other two domains (Fig. 5). The translation rate for eukaryotic genomes is approximately 3~8 amino acids per second (Mathews et al. 2000), compared to 10~20 amino acids per second for bacterial genomes (Liang et al. 2000). In contrast, eukaryotes have much larger genomes and, hence, many more proteins. Thus, eukaryotic proteins would undergo stricter translation selective pressure. This higher pressure may be one of the reasons for the higher co-adaption observed in eukaryotes. Focusing on the bacterial domain, we observed that larger bacterial genomes tended to have higher TAAIs, and the TAAI value correlated positively with the genome size. Higher selective pressure may be the reason for this positive correlation. Taking all of the results into account, co-adaption is one effect of translation selection at all three levels (domain, genome and protein) and the selection complies with the maximum efficiency & minimum cost principle.

**Fig. 5.**
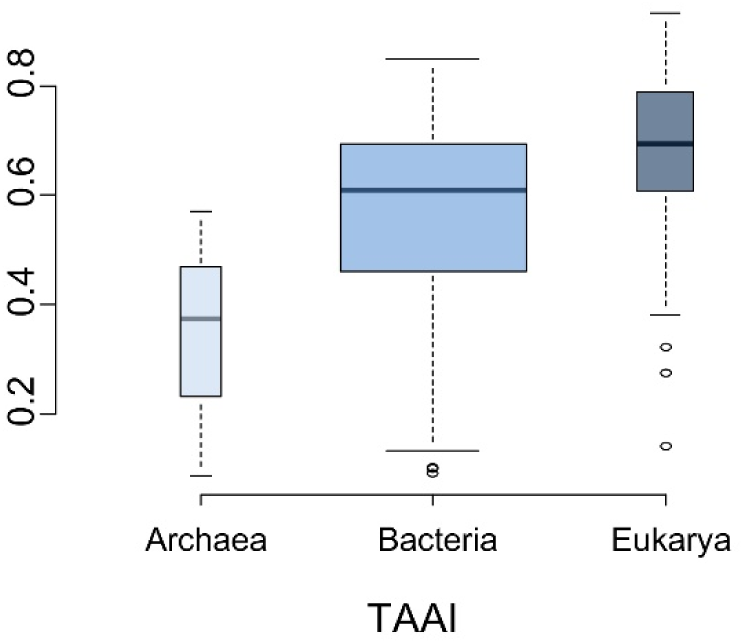
Comparison of the co-adaption (TAAI) for three domains. The medians for Archaea, Bacteria and Eukarya were 0.37, 0.61 and 0.69. Archaea had relatively low TAAIs, and Eukarya had the highest. Analysis o f variance revealed that the three domains were significantly different, with a p value close to zero (p = 2.57e-06).

This co-adaption can be applied to enhance translation efficiency in practice. Traditionally, in industrial application, the yield of a specific protein is improved in by optimizing its synonymous codon usage (Menzella 2011). A higher ratio of optimal codons could facilitate the transcription efficiency by frequent usage of abundant or efficient tRNAs (Cannarozzi et al. 2010). Here, the yield of EGFP was improved markedly in *E. coli* by optimizing TAAI through increasing the gene copy number of specific tRNAs, thus increasing the translation speed at least tenfold. This finding may be applied in industrial production. To obtain higher output of one protein, we could optimize its co-adaption between tRNA gene copy number and amino acid usage by adding specific tRNA gene copies. Thus, the protein’s translation could be accelerated quickly. One of the prominent advantages of such an operation is that the yields of multiple proteins could be improved in one round. The production of multiple proteins could be increased by adding specific tRNA gene copies corresponding to their amino acid usage. This ideal result is based on the supposition that adding a specific tRNA gene could increase the TAAIs of many proteins simultaneously. A more practical method would be to divide all target proteins into groups based on similar amino acid frequencies. If the tRNA genes to import are carefully chosen, the target group of proteins will have higher expression levels but the other proteins should remain almost unchanged.

In the field of synthetic biology, it is hoped to devise and construct a general bacterial chassis cell that integrates functional synthetic parts, devices and systems. In practice, such a chassis has often been constructed or synthesized based on small and slowly growing bacteria (Hutchison et al. 2016; Vickers 2016). However, slow growth may limit their capacity to produce enough target molecules in a short time. Our strategy of importing certain tRNA genes may help to address this problem when designing chassis cells.

## Methods

### *E. coli* strain and methods

DNA amplification and expression were performed in *E. coli* Top10 cells (F- *mcrA* Δ(*mrr-hsd*RMS-*mcr*BC) Φ80*lac*ZΔM15 Δ *lac*X74 *rec*A1 *ara*D139 Δ(*araleu*)7697 *gal*U *gal*K *rps*L (StrR) *end*A1 *nup*G). All bacterial media and methods used in this study were as described in Current Protocols in Molecular Biology (Ausubel et al. 1987).

### Production of synthetic genes

Oligonucleotides were synthesized using PCR amplification. The fragments were recombined to generate the target coding sequences, which were inserted behind the arabinose promoter. Positive clones were screened by resistance screening and confirmed by PCR and sequencing.

### Detection of target polypeptides

Cells were grown overnight at 30℃ in Luria-Bertani (LB) culture medium, and were inoculated in LB culture medium with ampicillin at OD600 = 2. After hours of constant shaking (OD600 = 0.6), L-arabinose (0.05%) was added to the culture medium to induce heterologous expression. Samples were collected at different time points and put on ice. When all samples were prepared, aliquots of the cells were observed through confocal microscopy (Leica TCS SP8, Germany), and the rest were collected by centrifugation (4000 x g, 20 min). The cells were then washed three times with cold PB (4000 x g, 10 min), and cell lysis buffer (phenylmethylsulfonyl fluoride [PMSF] 0.1 mM, PB 10 mM, lysozyme 1 mg/ml) was added to lyse the cells for 15min before sonication (3 min). After ultrasonic breakage, the samples were centrifuged, and the supernatants were collected. EGFP in the supernatants was measured using fluorospectro-photometer (Hitachi F7000, EX WL: 460.0 nm) and was quantified using a Bicinchoninic Acid (BCA) Protein Assay Kit (CWBIO) before adding loading buffer and boiling for five minutes. Thirty micrograms of the protein extract samples were loaded on 12% stain-free SDS-polyacrylamide gel (Bio-Rad), subjected to electrophoresis, and transferred to 0.2 µm polyvinylidene difluoride membranes (Millipore). After blocking with 5% nonfat milk in Tris-buffered saline buffer with Tween 20 (TBS/T) for 4h at room temperature, the membrane was incubated with the appropriate primary antibodies (Abmart GFP-tag mAb, 1:1000) for 18h at 4°C. Next, horseradish peroxidase (HRP)-conjugated goat anti-mouse secondary antibody (1:5000, ZSGB-BIO) was applied for 2h at room temperature. Finally, the signals were visualized using an Enhanced Chemiluminescence (ECL) kit (Roche).

## TAAI

TAAI is short for the “tRNA gene copy number and amino acid usage” accordance index and measures the co-adaption of amino acid usage and tRNA gene copy number. For a given protein sequence, the frequencies of the 20 types of amino acids are unique. For a specific organism, the average frequencies of amino acid usage also differ among organisms.

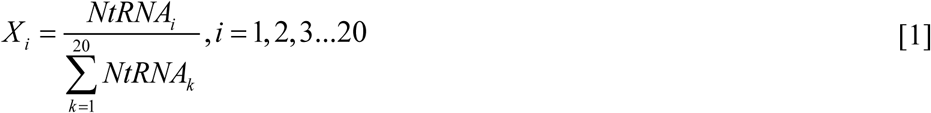

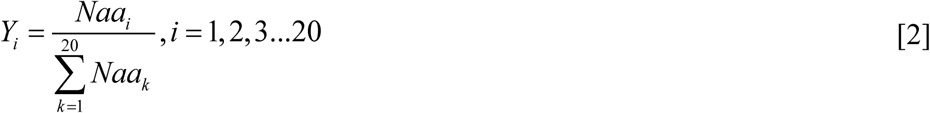

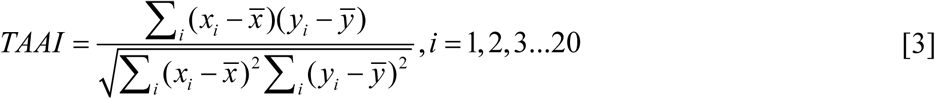

where Xi is the frequency of the gene copy of tRNA i in one organism’s genome in equation [1], and let Yi be the frequency of amino acid i of in a specific protein in the equation [2]. The value NtRNAi is the corresponding gene copy number of tRNAi decoding all codons for the ith amino acid. The value Naai is the corresponding counts of the ith amino acid in special protein or genome. TAAI in equation [3] is the Spearman correlation index of X and Y. In addition to the TAAI of one protein, we also calculated the general TAAI of an organism to denote its overall co-adaption. The overall value equals the average of all proteins’ TAAIs for that organism.

Where X_i_ is the frequency of the gene copy of tRNA i in one organism's genome in equation [1], and let Yi be the frequency of amino acid i of in a specific protein in the equation [2]. The value NtRNAi is the corresponding gene copy number of tRNAi decoding all codons for the ith amino acid. The value Naai is the corresponding counts of the ith amino acid in special protein or genome. TAAI in equation [3] is the Spearman correlation index of X and Y. In addition to the TAAI of one protein, we also calculated the general TAAI of an organism to denote its overall co-adaption. The overall value equals the average of all proteins' TAAIs for that organism.

## Bioinformatics data source

This work requires tRNA gene annotation information that is as accurate as possible; therefore, we compared annotation information from three databases. We chose three widely used databases: GenBank (Benson et al. 2013), a comprehensive bioinformatics database; the Genomic tRNA Database (Chan and Lowe 2009), which uses tRNAscan-SE (Lowe and Eddy 1997); and tRNAdb (Jühling et al. 2009), which contains more than 12,000 tRNA genes. A total of 410 genomes have the same tRNA gene annotation information in all three databases. If the tRNA gene annotations for one organism were consistent in the three databases, we took it as reliable and then employed this organism in further analysis. Protein sequences were acquired from GenBank, and protein abundance values were acquired from PaxDb (Wang et al. 2012).

We employed 53 organisms’ growth times (Table S4). Here, we chose those archaea and bacteria with growth times from (Rocha 2004). Among 410 organisms filtered for reliable tRNA gene annotation, 376 archaea and bacteria were chosen to perform the analysis of correlation between TAAI and genome sizes. When comparing the TAAIs among proteins within a specific genome, we analyzed the protein abundances and TAAIs in six model organisms, which have integrated abundances in PaxDb (Wang et al. 2012).

## Acknowledgments

We thank Mr. Lu-Wen Ning for suggestions, Juan Feng’s lab and Lixia Tang’s lab for providing help with experimental materials and methods, and our lab members for fruitful discussion. This work was supported by the National Natural Science Foundation of China [31470068 and 31501063] and the Fundamental Research Funds for the Central Universities of China [ZYGX2015J144].

